# Nanopore native RNA sequencing of a human poly(A) transcriptome

**DOI:** 10.1101/459529

**Authors:** Rachael E. Workman, Alison D. Tang, Paul S. Tang, Miten Jain, John R. Tyson, Philip C. Zuzarte, Timothy Gilpatrick, Roham Razaghi, Joshua Quick, Norah Sadowski, Nadine Holmes, Jaqueline Goes de Jesus, Karen L. Jones, Terrance P. Snutch, Nicholas Loman, Benedict Paten, Matthew Loose, Jared T. Simpson, Hugh E. Olsen, Angela N. Brooks, Mark Akeson, Winston Timp

## Abstract

High throughput cDNA sequencing technologies have dramatically advanced our understanding of transcriptome complexity and regulation. However, these methods lose information contained in biological RNA because the copied reads are often short and because modifications are not carried forward in cDNA. We address these limitations using a native poly(A) RNA sequencing strategy developed by Oxford Nanopore Technologies (ONT). Our study focused on poly(A) RNA from the human cell line GM12878, generating 9.9 million aligned sequence reads. These native RNA reads had an aligned N50 length of 1294 bases, and a maximum aligned length of over 21,000 bases. A total of 78,199 high-confidence isoforms were identified by combining long nanopore reads with short higher accuracy Illumina reads. We describe strategies for assessing 3′ poly(A) tail length, base modifications and transcript haplotypes from nanopore RNA data. Together, these nanopore-based techniques are poised to deliver new insights into RNA biology.

**DISCLOSURES:** MA holds shares in Oxford Nanopore Technologies (ONT). MA is a paid consultant to ONT. REW, WT, TG, JRT, JQ, NJL, JTS, NS, AB, MA, HEO, MJ, and ML received reimbursement for travel, accommodation and conference fees to speak at events organised by ONT. NL has received an honorarium to speak at an ONT company meeting. WT has two patents (8,748,091 and 8,394,584) licensed to Oxford Nanopore. JTS, ML and MA received research funding from ONT.

## INTRODUCTION

The roles of RNA in cell function are numerous and complex. Beyond the fundamental importance of mRNA, tRNA, and ribosomal RNA in translation, several classes of non-coding RNA (ncRNA) regulate cellular processes including division, differentiation, and programmed cell death^1^.

Sequencing by synthesis (SBS) strategies have dominated RNA sequencing since the early 1990s^2^. Typically this involves generation of cDNA templates by reverse transcription (RT)3,4 coupled with PCR amplification^5^. Sequential base identification along template strands is generated by DNA polymerase-dependent incorporation of complementary nucleotides into daughter strands. A high throughput version of this basic technique (RNA-seq6-12) can be implemented on a variety of platforms to determine both reference-based and *de novo* transcriptomes at high coverage^13^. RNA-seq has had an enormous impact in basic science and medicine exemplified by detailed maps of tissue-specific expression^14^, and by recent breakthroughs in classification of human cancers^15^. A single molecule SBS platform developed by Pacific Biosciences is used for reading long cDNA molecules end-to-end^16^. The Helicos platform was the first single molecule direct RNA sequencing platform which proved useful for counting transcripts at low RNA input^17^ but had limited utility due to its short read lengths.

Nanopore RNA strand sequencing has emerged as an alternative single molecule strategy^18-20^. It differs from SBS-based platforms in that native RNA nucleotides, rather than copied DNA nucleotides, are identified as they thread through and touch a nanoscale sensor. Nanopore RNA strand sequencing shares the core features of nanopore DNA sequencing, *i.e.* a processive helicase motor regulates movement of a bound polynucleotide driven through a protein pore by an applied voltage. As the polynucleotide advances through the pore in single nucleotide steps, ionic current impedance reports on the segment of bases that occupy a narrow reading head as a function of time. This series of ionic current segments is then used to infer nucleotide sequence using an algorithm trained with known RNA molecules.

Here we describe sequencing and analysis of a human poly(A) transcriptome from the GM12878 cell line using the Oxford Nanopore (ONT) platform. We demonstrate that long native RNA reads allow for discovery and characterization of RNA isoforms that are difficult to observe using short read cDNA methods^21,22^. Because native RNA strands are directly read by nanopores, nucleotide modifications and 3′ poly(A) tail lengths can be determined directly from the ionic current signal absent additional processing steps. Data and Resources are posted online at:

(https://github.com/nanopore-wgs-consortium/NA12878/blob/master/RNA.md).

## RESULTS

### RNA preparation, nanopore sequencing, and computational pipeline

The strategy we used to isolate and sequence native poly(A) RNA is shown in Figure 1a. Details are presented in the Online Methods. Briefly, total RNA was isolated from immortalized human B-lymphocyte cells (GM12878) using TRI Reagent Solution (ThermoFisher), followed by bead-based poly(A) selection. Approximately 750 ng of the poly(A) isolate was adapted for nanopore sequencing. This involved: i) attaching proprietary ONT adapters to the poly(A) RNA using T4 ligase; ii) generating poly(A) RNA/DNA duplexes by reverse transcription (RT); iii) ligating the adapted poly(A) RNA strand to a second proprietary ONT adapter bearing the RNA motor protein; and iv) loading the adapted poly(A) RNA onto individual MinION flow cells for sequencing using a standard ONT protocol (v). Generation of the cDNA complementary strand is not required, however we performed that step because it is reported to improve throughput^18^. A nanopore ionic current trace for TP53 mRNA (Figure 1b) shows typical features for poly(A) RNA translocation.

**Figure 1.**
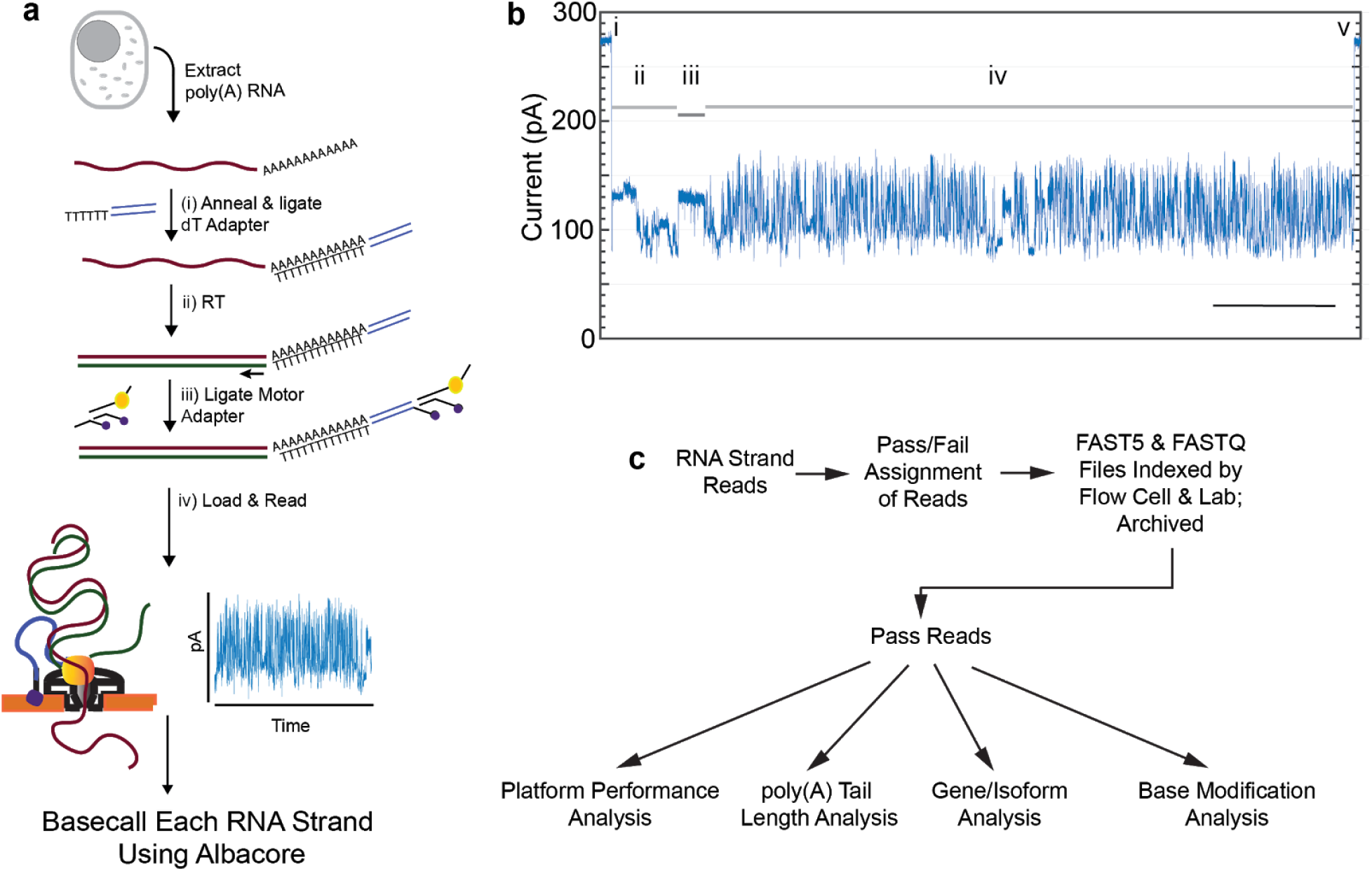
Nanopore native poly(A) RNA sequencing pipeline. (a) RNA is isolated from cells followed by poly(A) selection using poly(dT) beads. poly(A) is then prepared for nanopore sequencing using the following steps: (i) A duplex adapter bearing a poly(dT) overhang is annealed to the RNA poly(A) tail, followed by ligation of the strand abutting the poly(A) tail; ii) the poly(dT) complement is extended by reverse transcription; iii) a proprietary ONT adapter bearing a motor enzyme is ligated to the first adapter; and (iv) the product is loaded onto the ONT flow cell for reading by ionic current impedance. The ionic current trace for each poly(A) RNA strand is base called using a proprietary ONT algorithm (Albacore). (b) A representative ionic current trace for a 2.3 kb TP3 transcript ionic current components: (i) Strand capture; ii) ONT adapter translocation; iii) poly(A) RNA tail translocation; iv) mRNA translocation; and (v) exit of the strand into the trans compartment. Bar is 5 seconds (c) Processing of the RNA strand reads *in silico*, followed by data analysis.

The ionic current readout for each poly(A) RNA strand was basecalled using Albacore version 2.1.0 (ONT). The resulting sequences were then classified as either pass or fail based on a per-read average Phred-scale quality value threshold of 7. FASTQ and FAST5 files were indexed using nanopolish^23^ to associate individual sequences with their corresponding ionic current traces. We also performed nanopore cDNA sequencing of the GM12878 poly(A) RNA using the same RNA sample and analysis pipeline, but with modified parameters appropriate for cDNA sequencing (Online Methods). Both the RNA and cDNA data were archived and used for downstream analyses (Figure 1c). They are available on GitHub at:

(https://github.com/nanopore-wgs-consortium/NA12878/blob/master/RNA.md).

### Native poly(A) RNA sequencing statistics

Each of six consortium sites performed five separate nanopore sequencing runs. Together, these thirty runs produced 13.0 million poly(A) RNA strand reads, of which 10.3 million qualified as pass reads (Q-value threshold 7). Throughput varied substantially between 50K and 831K pass 1D poly(A) reads per MinION flow cell (median = 372K; S.D. = 260K). The 10.3 million pass 1D RNA nanopore reads had an N50 length of 1334 bases, and a median length of 771 bases (Table 1). Reads were aligned to the GRCh38 human genome reference sequence using minimap2 with a splice-aware setting (-ax splice-uf-k14)^24^. This algorithm was chosen because it can align nanopore reads to exons in the human genome while accurately spanning across introns^25^. Of the 10.3 million pass reads, 9.9 million (96.5%) aligned to the reference. The 360,000 unaligned pass reads had a median read length of 211 bases, which suggests that shorter nanopore reads were more difficult to align.

**Table 1.**
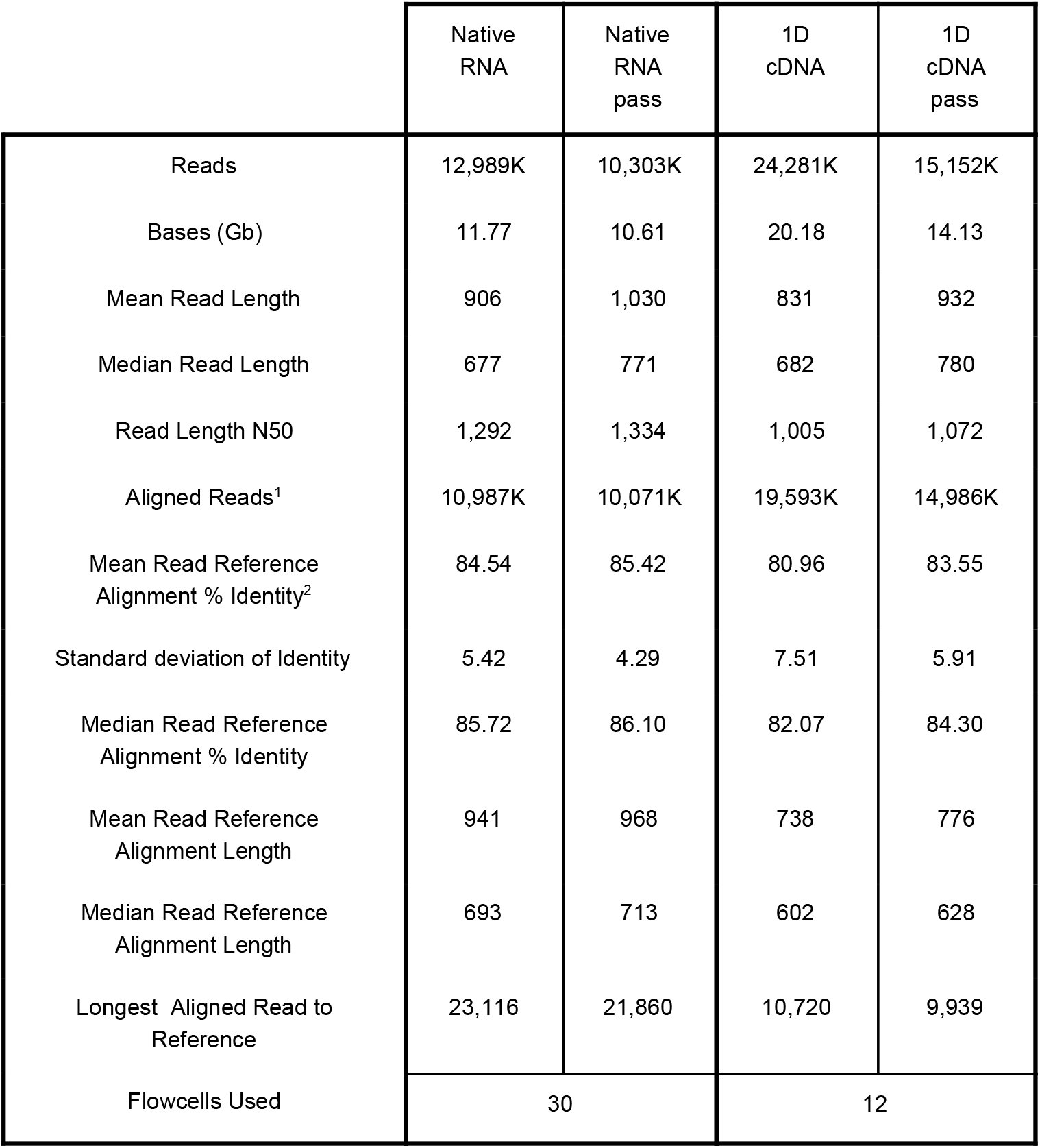
Yield and read alignment statistics for native RNA and 1D cDNA. Aligned reads (1) refers to reads aligned against GENCODE v27 transcripts using bwa-mem. Mean Read Reference Alignment % Identity (2) was calculated using scripts described in Quick *et al.^77^*. Pass reads indicate an Albacore generated q-score of 7 or greater.

We also aligned the RNA reads to a GRCh38 reference transcriptome (GENCODE v27) using minimap2^24^. The -ax map-ont setting is designed to align ONT reads to reference sequences with technology-specific mapping parameters. A comprehensive list of the genes and isoforms represented among the aligned native RNA reads can be found on GitHub and in **Supplementary Tables 1** and **2** respectively. MarginStats^26^ was employed to calculate the number of matches, mismatches, and indels per aligned read in this population. We found a median identity of 86% (Figure 2a), with mismatch, insertion, and deletion errors of 2.4%, 4.3%, and 4.4% respectively. Percent identity was consistent across institutions and among flow cells (median average identity of pass reads equal to 85.5%, with a standard deviation of 0.65).The basecaller seldom confused G-for-C or C-for-G (0.38% and 0.47% errors respectively); by comparison, C-to-T and T-C errors were substantially higher (3.62% and 2.23% respectively) (Figure 2b).

**Figure 2.**
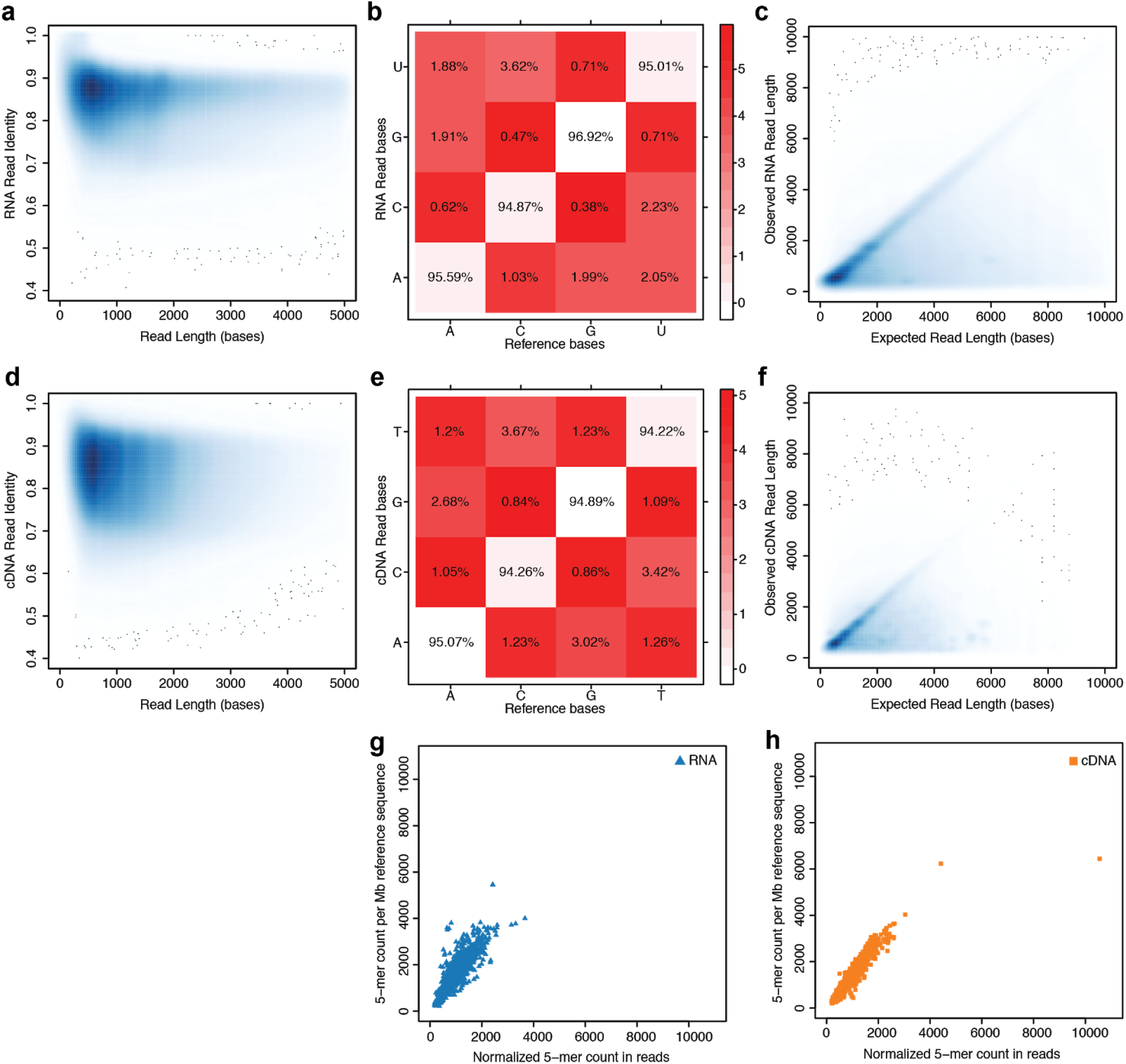
Performance statistics for nanopore native RNA and cDNA sequencing. (a) Alignment identity vs. read length for native RNA reads. (b) Substitution matrix for native RNA reads. (c) Observed v. expected read length for native RNA reads. (d) Alignment identity vs. read length for cDNA reads. (e) Substitution matrix for cDNA reads. (f) Observed vs. expected read length for cDNA reads. (g) Observed vs expected kmers (k = 5) for native RNA reads. and (h) Observed vs. expected kmers (k = 5) for cDNA reads.

Nanopore RNA reads that aligned to the GENCODE v27 transcriptome reference ranged from 85 nt (a fragment of an mRNA encoding Ribosomal Protein RPL39), to 22kb (an mRNA encoding spectrin repeat containing nuclear envelope protein 2 (SYNE2). RNA reads were also aligned to a set of high-confidence isoform sequences that were curated using a pipeline termed ‘FLAIR’ (see below). Using these alignments, we compared observed vs expected length and found general agreement (Figure 2c).

For nanopore cDNA data, we observed a median identity of 85% (Figure 2d) which is comparable to other recent nanopore DNA studies^27^. The substitution error patterns for cDNA data were similar to those for native RNA data (Figure 2e). However, while there was agreement in observed vs expected read lengths for cDNA and RNA, there were substantially fewer cDNA reads above 4 kb in length compared to the RNA reads (Figure 2c,f).

### Mitochondrially-encoded transcripts

Mitochondrially-encoded transcripts are essential and abundant in virtually all eukaryotic cells, and unlike most nuclear genes, they are each transcribed from a single exon. This simplifies measurement of RNA strand physical properties.

The human mitochondrial genome (MT-genome) is a closed circle composed of 16,569 nucleotides^28^ encoding 13 proteins involved in oxidative phosphorylation, 16S and 12S mitochondrial ribosomal RNAs (MT-RNR2 and MT-RNR1), and humanin (a short protein encoded within MT-RNR2^29^). All but MT-ND6 (NADH-ubiquinone oxidoreductase chain 6) are found on the heavy strand (H strand) of the circular duplex. There are also 22 mitochondrially-encoded tRNA genes, often situated between protein coding genes. Protein-encoding mRNAs are transcribed as long intact polycistronic molecules for both the H and L (light) strand. These are cut into individual gene-specific transcripts principally by enzyme-dependent excision of tRNAs^30^.

Of the 9.9 million aligned poly(A) RNA strand reads, 911,588 (*~*10%) aligned to the mitochondrial H strand and 39,291 reads (~0.4%) aligned to the L strand (Figure 3a). Mean coverage across the mitochondrial reference was ~6,600X. Every nucleotide position of the H strand was covered at least 7X; two positions of the L strand (positions 433 and 956 relative to the 16,569 nt MT-genome) were not covered (https://goo.gl/erWFyu).

**Figure 3.**
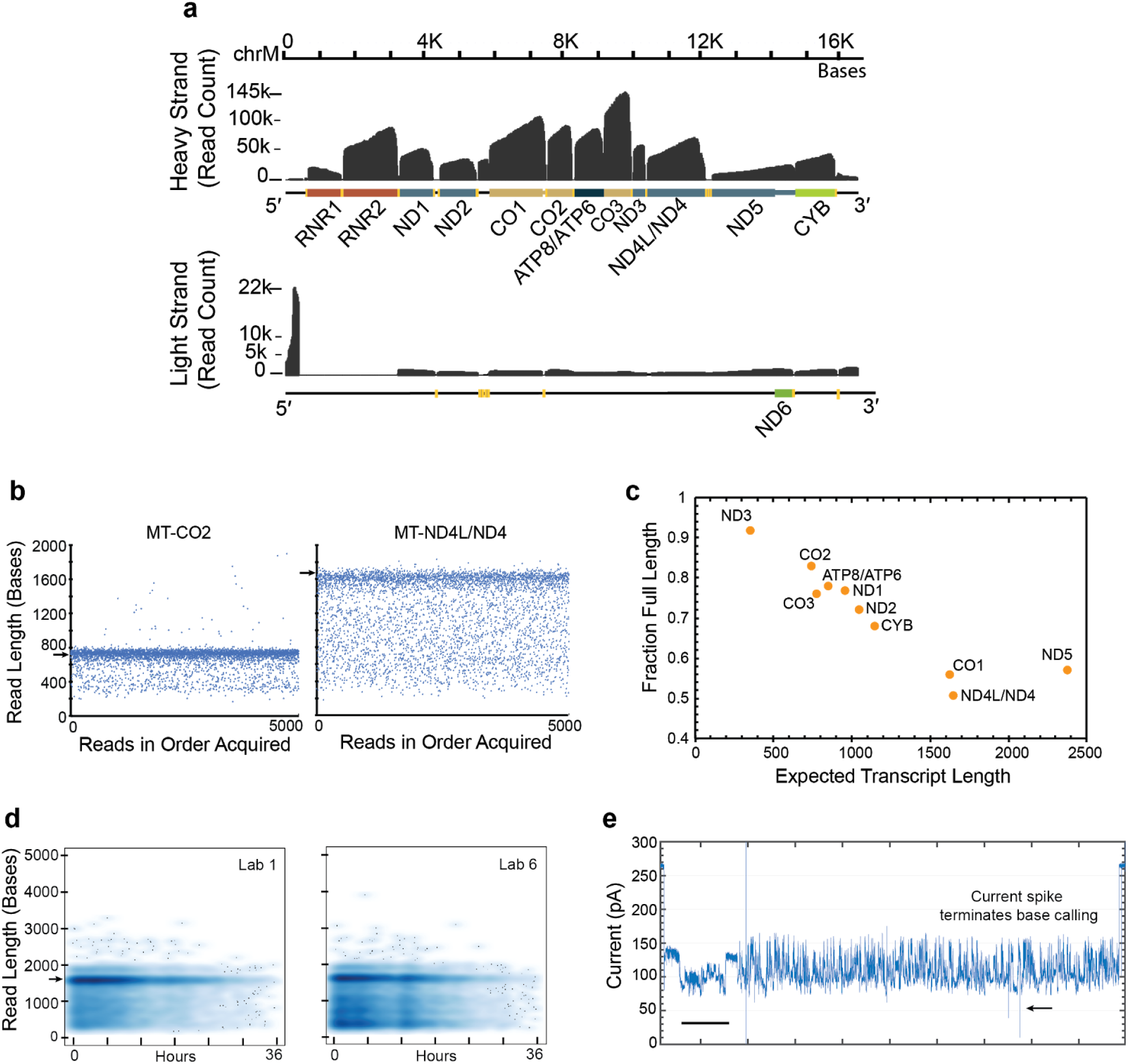
Mitochondrially-encoded poly(A) RNA transcripts. (a) Read coverage of the H strand (top) and the L strand (bottom). Dark grey is base coverage along the MT genome. Labelled colored bars represent protein coding genes including known UTRs, or ribosomal RNA (RNR1,RNR2). Texts denote specific genes absent the MT prefix. Yellow bars represent tRNA. (b) Distribution of nanopore read lengths for MT-CO2 and MT-ND4L/ND4 transcripts. Each point represents one of approximately 5000 reads in the order acquired from a single Lab 1 MinION experiment. Horizontal arrows are expected transcript read lengths. (c) Relationship between expected transcript read length and fraction of nanopore poly(A) RNA reads that were full length. Each point is for a protein coding transcript on the H strand. Labels are for mitochondrial genes absent the MT prefix. See Online Methods for definition of ‘Full Length’. (d) MT-CO1 poly(A) transcript read length vs MinION run time. The panel at left is from lab 1 and is representative of results from labs 2-5. The panel at right is from lab 6 which yielded results that differed substantially from the norm. (e) Ionic current trace for translocation of a MT-CO1 transcript. It is representative of traces wherein base calling was artificially truncated by a signal anomaly. The time bar is two seconds.

Overall, the nanopore RNA reads recapitulated established features of the human MT-transcriptome including: i) the tRNA punctuation model of MT-mRNA processing^30^; ii) 3′ UTR of MT-CO1, MT-CO2, and MT-ND5 mRNAs^31^; iii) bicistronic transcripts comprised of MT-ND4L/MT-ND4 and MT-ATP6/MT-ATP8; and iv) precursors such as RNA19 which is composed of uncleaved RNR2-TRNL1-ND1 segments. Nanopore sequencing also detected MT poly(A) RNA strands that are difficult to observe by conventional means. These included; i) an individual 9.1kb polycistronic L strand transcript that extended from within MT-TC (position 5,819, MT-genome) to within the ORF of ND6 (position 14,651, Mt-genome); ii) MT-CO1 transcripts bearing *Ori_L_* bases at their 5′ ends^31^; and iii) polycistronic reads bearing pre-processed copies of all 22 MT-tRNA.

Systematic analysis of H strand mRNA read lengths revealed strengths and limitations of the current ONT platform. Figure 3b compares 5,000 reads that aligned to MT-CO2 or to MT-ND4L/ND4 genes versus the order of read acquisition. In each panel, the dominant band corresponded closely to the expected transcript length (732 nt and 1,673 nt for MT-CO2 and MT-ND4L/ND4 respectively). These data also revealed a marked drop in the number of reads below 300 nt, and virtually no reads below 200 nt. This suggests that short RNA reads cannot be sequenced using the present ONT software.

A population of poly(A) reads appeared randomly distributed between the dominant band and about 300 nt for both MT-CO2 and MT-ND4L/ND4. We reasoned that if random, these apparent truncations would be equally probable along the length of a given poly(A) RNA strand. If so, the proportion of full length reads should decrease linearly as a function of expected transcript read length. To test this reasoning, we quantified the fraction of full length reads for protein-coding transcripts of the mitochondrial H strand (Online Methods) and found a strong linear anti-correlation in most cases (Figure 3c). The single outlier was MT-ND5, the mitochondrial transcript with a 568 nt 3′ UTR.

H strand poly(A) RNA truncations could occur at any of several non-biological steps during the sequencing process, or they could arise from regulated enzymatic degradation in the mitochondrion^32^. We first considered non-biological causes of truncation. One possibility was strand breaks during sequencing on the nanopore flow cell, which takes place at 34°C in the presence of Mg^2+^. As a test, we analyzed MT-CO1 read length distribution for each of the six laboratories as a function of time on the ONT flow cells. In Figure 3d, results for Lab 1 (left panel) are representative of results for five of the six laboratories. A band consistent with full length MT-CO1 is evident, as is a diffuse distribution of shorter reads. The number of reads of all lengths declined steadily over 36 hours, consistent with declining throughput with time previously described for ONT devices. Importantly, the fraction of full length reads declined only slightly (5%). The results for Lab 6 (Figure 3d, right panel) were similar in many respects to those for Labs 1-5, including the modest decline in full length read fraction with time (5%). However, the proportion of truncated reads at the beginning of the experiment was much higher for Lab 6. This underscores the importance of RNA sample quality during preparative steps.

Truncation of RNA reads could also be caused by physical nanopore sequencing artefacts such as enzyme stalls during RNA translocation, or extraneous voltage spikes that convolute the ionic current signal. As an initial test to establish the source of these truncations, we curated 101 random MT-CO1 ionic current traces acquired by Lab 1. We found that 48 of the curated traces were full length or longer, consistent with comprehensive data for MT-CO1 (Figure 3c). Thirty-five had ionic current traces that appeared normal except for shorter translocation times through the nanopore, and associated shorter sequence lengths. The remainder (18 traces) had measurably shorter sequence lengths that were truncated by anomalous, noisy prolonged ionic current patterns, or by voltage spikes that correlated with termination of base calling in the time domain (Figure 3e). We conclude that approximately 20% of the MT-CO1 poly(A) RNA reads were truncated by nanopore signal noise.

### Isoform detection and analysis

We reasoned that long nanopore reads could improve our ability to resolve RNA exon-exon connectivity, allowing for discovery of novel RNA isoforms and more accurate quantification of RNA transcript levels. However, individual nanopore reads average 14% per-read error rates, confounding the precise determination of splice sites. To overcome this limitation, we generated a subset of filtered and corrected nanopore reads from the total poly(A) RNA population and then identified high-confidence isoforms using FLAIR^25^ (Full-Length Alternative Isoform Analysis of RNA, Online Methods).

This strategy is summarized as follows. We first corrected the splice site boundaries of the nanopore poly(A) RNA read alignments to the genome using short-read Illumina cDNA data for GM12878 and existing transcript annotations from GENCODE v24. After correction, we saw marked improvement in splice site accuracy (**Supplementary Fig. 1**). Given the observed truncations in nanopore poly(A) reads, we wanted to increase the confidence of transcription start sites (TSSs); therefore, we considered only reads with 5′ ends proximal to promoter regions as defined by ENCODE promoter chromatin states from the GM12878 cell line^33-35^. We then used FLAIR to define isoforms specific to GM12878 in two steps: i) reads were grouped into isoforms according to the exact splice junctions used in genomic alignments; and ii) only isoforms supported by at least five reads were retained.

This analysis resulted in 78,199 high-confidence isoforms representing 10,513 genes, henceforth referred to as isoform Set A. In Set A, for genes with assembled isoforms, a majority (62.6%) contained at least one previously annotated isoform and at least one novel isoform. An example set of novel isoforms arose from a novel transcription start site with multiple splice variants for a lncRNA on chromosome 17 (Figure 4a). We also generated a more stringent population (Set B) by filtering for isoforms with unique splice junction chains from set A, thus removing isoforms that could be truncated transcripts of longer ones. Set B contains 51,039 isoforms and was used in all further analyses. Defining novel isoforms as those which contain a novel set of splice junctions not found in GENCODE v24, 65.3% of isoforms in Set B are novel, comprising 30.2% of total reads.

**Figure 4.**
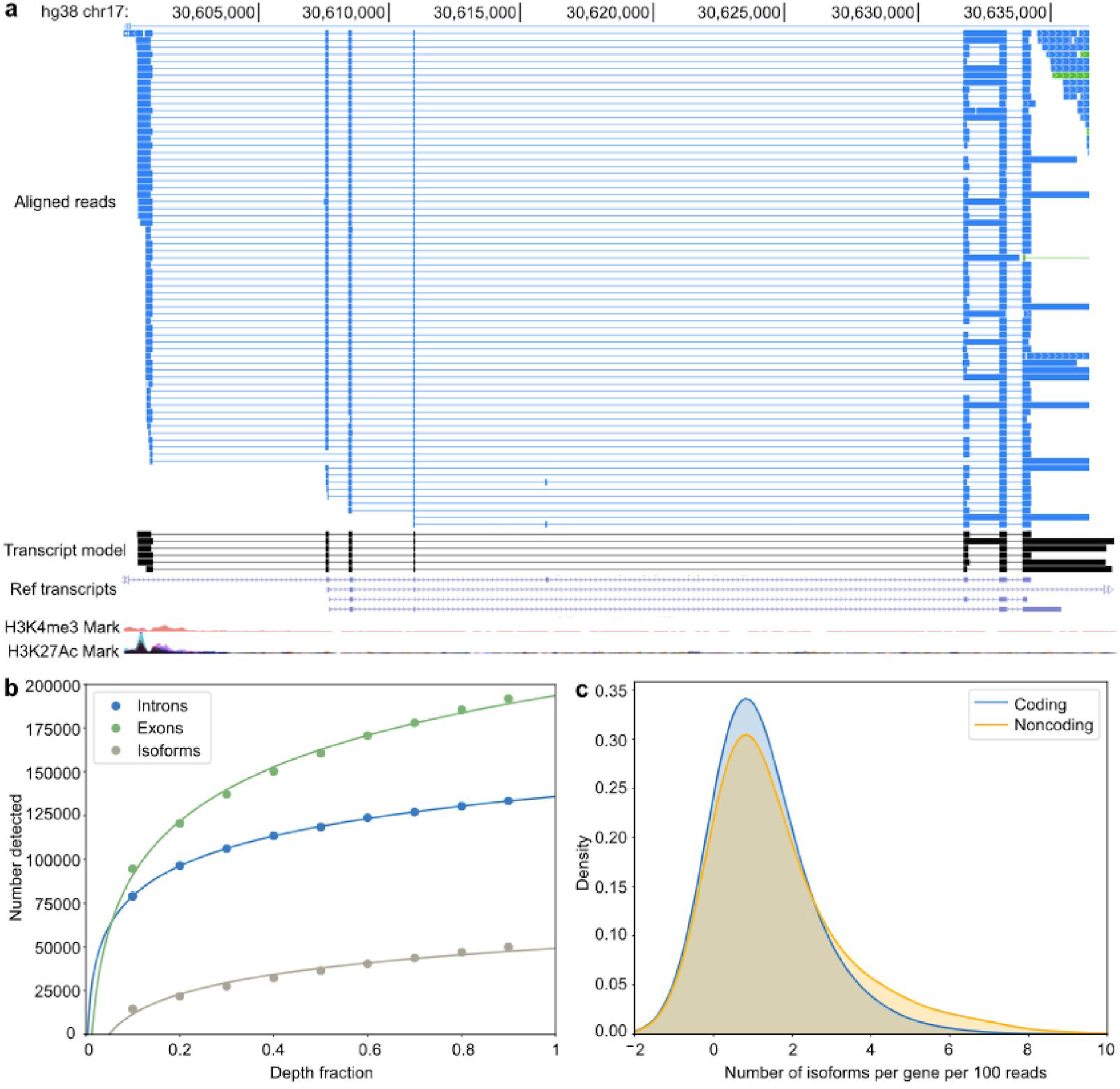
Isoform-level analysis of GM12878 native poly(A) RNA sequence reads. (a) Genome browser view of novel isoforms found in the native RNA data. From top to bottom, the tracks are: a subset of the aligned native RNA reads; the FLAIR-defined isoforms; Gencode gene annotation; transcription regulatory histone methylation marks; and transcription regulatory histone acetylation marks. (b) Saturation plot showing the number of introns, exons, and isoforms discovered (y-axis) in relation to the different fractions (x-axis) of total native RNA data used for FLAIR isoform definition. (c) Distributions of the number of isoforms per gene categorized as coding or noncoding, normalized per 100 aligning reads to the gene.

To determine if the ~9.9 million aligned native RNA sequence reads was sufficient to reach saturation of discovered isoforms, we subsampled the native RNA reads in increments of 10% and defined isoforms using the subsampled datasets. As expected, increasing sequencing depth increases the number of isoforms discovered in a nonlinear manner (Figure 4b).

We segregated the high-confidence isoforms into three categories: i) long non-coding isoforms (lncRNAs) that lacked an annotated start codon; ii) unproductive isoforms that contained ORFs but with premature termination codons upstream of the last splice junction; and iii) protein coding isoforms (Online Methods). When combined, noncoding isoforms were more numerous than protein coding isoforms per gene shown in Figure 4c (Wilcoxon p=3.72e-5). This finding is consistent with previous studies demonstrating increased levels of alternative splicing in noncoding exons^36,37^.

### Kmer coverage

Kmer coverage analysis has proved useful for identifying DNA segments that are difficult to sequence using the ONT MinION^26,27^. Historically, problematic DNA kmers often contained short homopolymer stretches^26^. Implementing a similar comprehensive kmer analysis for transcriptomes would be useful as well, however it is not straightforward because RNA expression levels inherently vary between genes, and because the composition of isoforms expressed at low levels can be difficult to characterize unambiguously.

As a first approximation, we assessed nanopore RNA kmer coverage using a set of high-confidence full-length RNA isoforms as the sample sequences. Briefly, FLAIR isoforms from set B were filtered for those corresponding to known 5′ start and 3′ stop sites in expressed gene isoforms (Online Methods). With this high-confidence mapping of individual reads to genomic sequences we determined RNA kmer frequencies that took into account numbers of sequence reads for individual isoforms and based expected kmer content on the genomic (DNA) sequences corresponding to each high-confidence full-length isoform. Of the 10.3 million pass reads, 8.2 million reads were each assigned to a high-confidence isoform. For comparison, of 15.1 million pass cDNA reads, 10.2 million pass cDNA reads were each assigned to a high-confidence isoform. These reads included all possible kmers which were represented in sufficient numbers to permit a statistically valid analysis. Figure 2g and 2h show normalized counts for all 1024 kmers determined from native RNA and cDNA data respectively. Kmer frequencies exhibit an approximate one-to-one trend between observed and expected frequencies for both native RNA and cDNA. The largest deviation often occurred for homopolymer-rich kmers that were underrepresented in native RNA and overrepresented in cDNA (**Supplementary Tables 3** and **4** respectively).

### Assignment of transcripts to parental alleles using nanopore reads

The long reads produced by nanopore RNA sequencing should in theory be easier to assign to parental allele of origin, due to the greater chance of encountering a heterozygous SNP. Using HapCUT2^38^, we assigned transcript reads which contained at least two heterozygous variants to their parental allele of origin. This pipeline found 2,917 genes with at least 10 haplotype informative reads: 2,885 autosomal genes and 32 from the X chromosome (**Supplementary Table 5**). Within the autosomal genes, 464 (16%) showed significant allele specific expression (ASE) (binomial test, P < 0.01), while the remainder demonstrated balanced expression from both alleles. Conversely, when we focused solely on genes from the X-chromosome, we observe clear skewing of expression consistent with X-inactivation. Of the X-chromosome genes with allele-informative SNVs, 28 (88%) showed significant ASE (binomial test, P < 0.01); and of the maternal genes with determined ASE, 26 (93%) showed expression from the maternal allele. The only reads aligning to the paternal X-chromosome map to the genomic region for Xist (Figure 5a), a lncRNA known to be expressed from the inactive X-chromosome and recruit epigenetic silencing machinery^39^. Greater than five heterozygous variants at the Xist locus in GM12878 were used to confirm paternal expression of Xist, two such variants are highlighted in Figure 5a.

**Figure 5.**
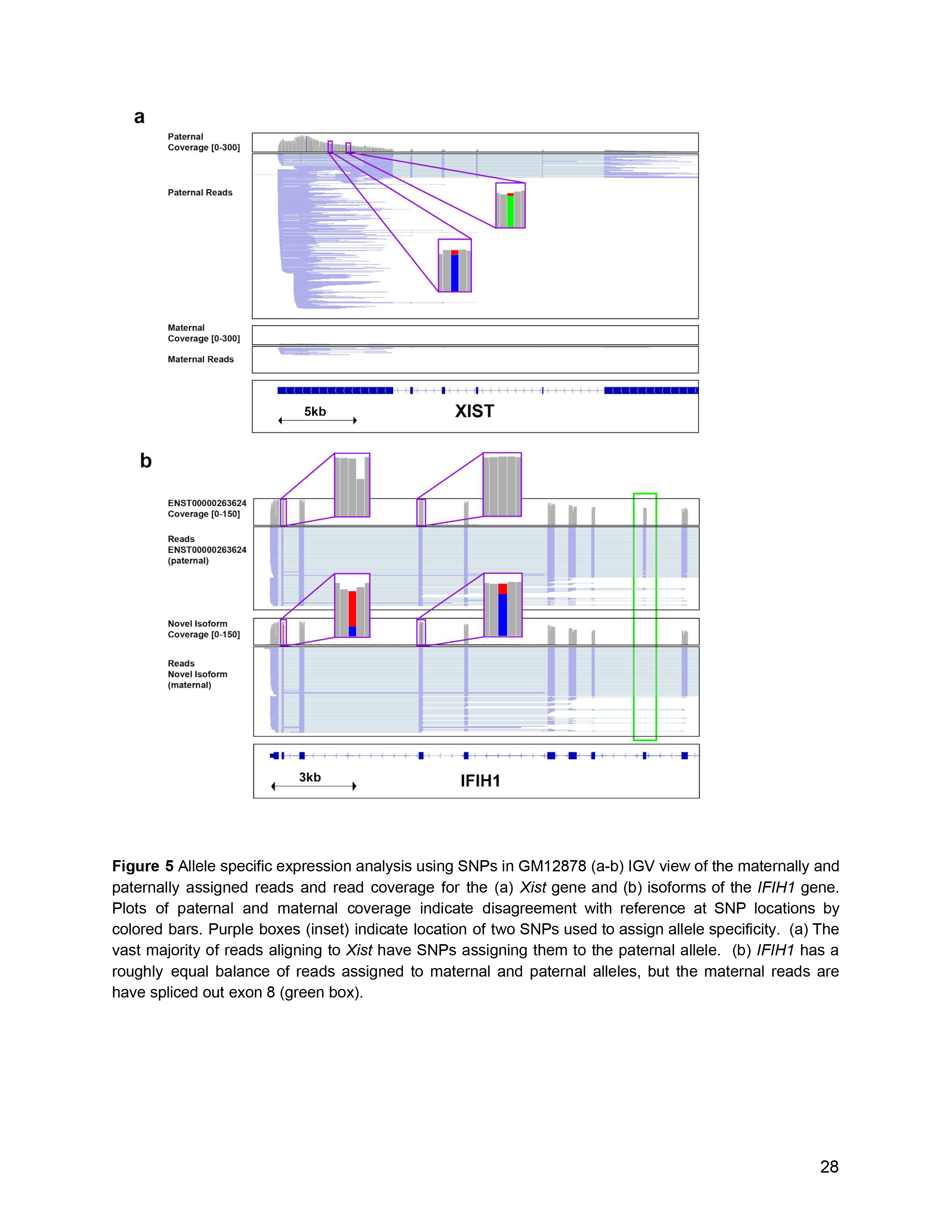
Allele specific expression analysis using SNPs in GM12878 (a-b) IGV view of the maternally and paternally assigned reads and read coverage for the (a) *Xist* gene and (b) isoforms of the *IFIH1* gene. Plots of paternal and maternal coverage indicate disagreement with reference at SNP locations by colored bars. Purple boxes (inset) indicate location of two SNPs used to assign allele specificity. (a) The vast majority of reads aligning to *Xist* have SNPs assigning them to the paternal allele. (b) *IFIH1* has a roughly equal balance of reads assigned to maternal and paternal alleles, but the maternal reads are have spliced out exon 8 (green box).

We then combined our allele-specific reads with isoform assignments to mine our data for allele-specific isoforms (Online Methods). We identified 34 genes with discordant allele specificity in two isoforms, i.e. >80% of reads expressed from only one allele in one isoform, and >80% of reads from the other allele in another isoform (**Supplementary Table 6**). One of these genes, *IFIH1*, has a paternal isoform with exon 8 retained, while the maternal isoform excluded exon 8 (Figure 5b). Further, the closest SNV used in allele-assignment is 886nt away from the alternate splicing event in the transcript, making the allele-specificity of this novel *IFIH1* isoform difficult or impossible to detect with short read sequencing.

### 3′ poly(A) analysis

The ionic current signal arising from native poly(A) RNA strand translocation contains a distinct low variance component near the 3′ end that is attributable to the poly(A) tail (Figure 1b,iii). We have developed a computational method (‘nanopolish-polya’) to segment this signal into four regions which allows us to estimate how many ionic current samples were drawn from the poly(A) tail region (**Supplementary Note 1**). Then, by correcting for the rate at which the RNA molecule passes through the pore, nanopolish-polya estimates the length of the poly(A) tail in nucleotides.

To test this method we obtained six poly(A) RNA control datasets generated by ONT. These datasets consisted of ionic current traces for synthetic *S. cerevisiae* enolase transcripts appended with 3′ poly(A) tails of 10, 15, 30, 60, 80 or 100 nucleotides (see ENA accession PRJEB28423 for details of how these were generated). There was a second version of the 60nt poly(A) tailed construct (60nt-N) which contained all possible combinations of a 10nt random sequence inserted between the enolase sequence and the 3′ poly(A).

Poly(A) tail length estimates for these synthetic controls are shown in Figure 6a, and statistics are reported in **Supplementary Table 7**. Median estimates fell within 4 nucleotides of the expected tail length for the 10-to-80 poly(A) datasets; for the 100nt dataset, the median estimate was 109nt. We observed that 66%-80% of the estimated lengths fell within 2 median absolute deviations of the expected tail length. The predicted tail length distribution for the 60nt-N data set (bearing the 10nt random sequence insert) contained a higher proportion of short poly-A tails than expected, which may indicate amplification errors specific to this sample.

**Figure 6.**
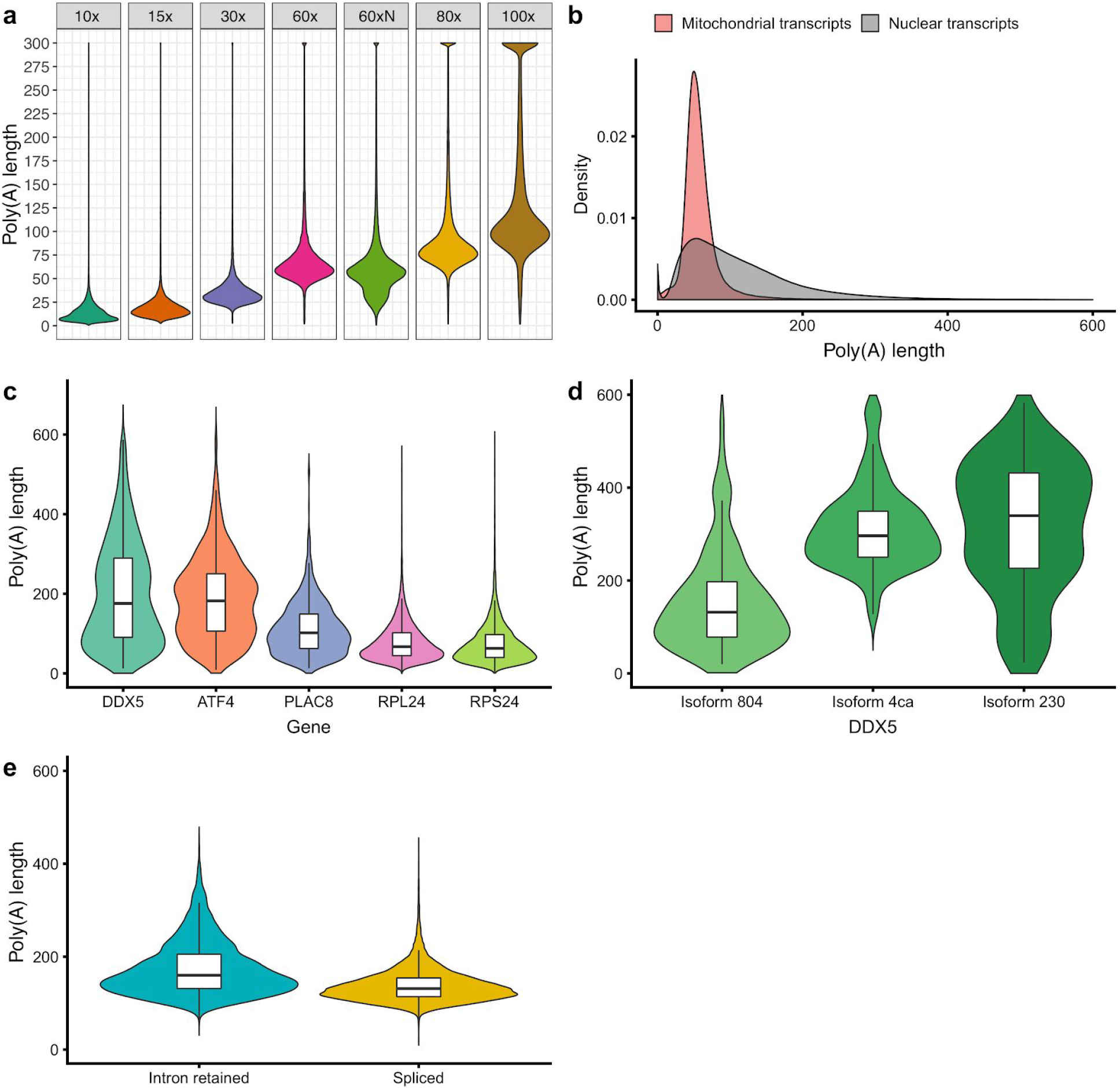
Testing and implementation of the poly(A) tail length estimator nanopolish-polya. (a) Estimate of poly(A) lengths for a synthetic enolase control transcript bearing 3′ poly(A) tails of 10, 15, 30, 60, 80 or 100 nucleotides. ‘60xN’ contained all possible combinations of a 10nt random sequence inserted between the enolase sequence and the 3′ poly(A) 60mer. (b) Poly(A) length distributions for transcripts encoded in the mitochondrial genome versus nuclear-encoded genes. (c) Violin plots showing the range of poly(A) tail lengths sequenced, with the longest (DDX5, ATF4), shortest (RPL24, RPS24), and average (PLAC8) poly(A) distributions plotted. (d) Distribution of poly(A) tail lengths for three isoforms of DDX5 plotted. (e) Distribution of poly(A) tail lengths for spliced and intron retaining transcripts identified in FLAIR isoform Set B.

A limitation of our approach is the inability to detect when the poly(A) region stalls in the nanopore sensor during translocation. When this occurs the program may overestimate the length of the poly(A) tail, with the degree of overestimation dependent on the duration of the stall. From the control data, we estimate this occurs for 1-3% of the sequenced molecules. As the length of the poly(A) tail increases, the variance of our estimator does as well. This is expected because the number of samples in the tail region for a fixed expected tail length is not deterministic and has substantial variation due to the kinetics of translocation. Heuristically, we observed that the number of samples corresponding to a single sequenced nucleotide can be modelled as a Gamma-distributed random value. If we approximate the number of samples in the poly(A) tail as a sum of independent random values, we expect the variance to increase commensurately with the tail length. We found that we were able to offset some of this inherent variance by using the overall transcript translocation rate as an estimator of the poly(A) rate (further details are provided in the **Supplementary Note 1**).

We applied this poly(A) length estimator to the complete GM12878 native poly(A) RNA sequence dataset. Overall, the poly(A) length distribution centered at approximately 50nt, with a broad dispersion of longer poly(A) tails for some transcripts (Figure 6b). When we segregated mitochondrial-encoded transcripts from nuclear-encoded transcripts, we found that the mitochondrial transcripts had poly(A) lengths which peaked at 52nt, with a mean of 59 and almost no poly(A) tail lengths greater than 100nt (Figure 6b). This is consistent with results for mitochondrial poly(A) RNA from other human cell lines^31^. Conversely, nuclear transcripts showed a much larger size distribution, with the distribution peaking at 58nt, with a mean of 112nt and a large number of poly (A) tails greater than 200nt.

Next we examined poly(A) tail length differences between genes and between isoforms of individual genes. To avoid biases associated with small sample sizes, we analyzed only high coverage genes. We divided the datasets into three bins: 100-1,000 reads per gene, 1,000-20,000 reads per gene, and greater than 20,000 reads per gene. Within the 1,000-20,000 dataset, numerous examples were identified where genes had poly(A) lengths that differed greatly from the mode (Figure 6c). These included the RNA-binding splicing protein, DDX5, and the CREB-family transcription factor, ATF4. Within violin plots for these genes, multiple poly(A) length peaks were observed, suggesting numerous poly(A) tail sub-populations. The short median poly(A) length for ribosomal proteins (RPL24, RPS24) is largely in agreement with the published literature^40^ (Figure 6c).

We also found that isoforms for a given gene identified by our FLAIR analysis could have different poly(A) tail lengths. For example, within the highly expressed DDX5 gene, different isoform transcripts contained measurably different poly(A) lengths (Figure 6d, **Supplementary Figure 2**). Notably, a previously annotated isoform (ENST00000581230; ‘230’) had a poly(A) tail peak mode of 450nt, and a newly identified isoform (Isoform 4ca, this study) had a poly(A) tail peak mode of 300nt. Both of these isoforms were incompletely processed, with some retained introns. By comparison, the remainder of the DDX5 isoforms had poly(A) lengths centered below 100nt.

The difference in the isoforms of DDX5 motivated us to explore the relationship between poly(A) length and RNA splicing. For this analysis, we classified each isoform in FLAIR Set B as either completely spliced or incompletely spliced (i.e. intron-retaining). We observed that transcripts with retained introns tended to have longer poly(A) tails (median 161nt) than did spliced transcripts (median 132nt) (Figure 6e). This result is consistent with a previous observation that nuclear transcripts with retained introns have longer poly(A) tails due to the activity of the nuclear poly(A) binding protein (PABPN1) and poly(A) polymerase (PAP)^41^.

### Modification detection

Nucleotide modifications can affect RNA shape, local charge and base-pairing potential, thus altering protein-binding affinity^42^. N6-methyladenine (m6A) is the most common internal modification on mRNA^43^, and has been implicated in many facets of RNA metabolism^44^. m6A dysregulation has been linked to deleterious effects on cell differentiation and stress tolerance, and to a number of human diseases, including obesity and cancer^45^. Because m6A modifications are enriched in 3′ UTRs, the impact of this modification appears to be largely regulatory, as opposed to altering protein coding sequence. Additionally, it has been reported that roughly two-thirds of 3′ UTR regions with m6A modifications contain miRNA sites, indicating that these features could be related^46^.

We used existing immunoprecipitation studies to identify a list of high abundance genes likely to contain m6A^47,48^ in the nanopore poly(A) RNA dataset. We focused our studies on the GGACU binding motif of METTL3, a subunit of the m6A methyltransferase complex^47^. As an example, we compared the raw current signal at putative m6A sites for Eukaryotic elongation factor 2 (*EEF2)* RNA versus the signal for an *in vitro* transcribed copy produced from GM12878 mRNA (Online Methods). This comparison revealed an ionic current change attributable to m6A (Figure 7a).

**Figure 7.**
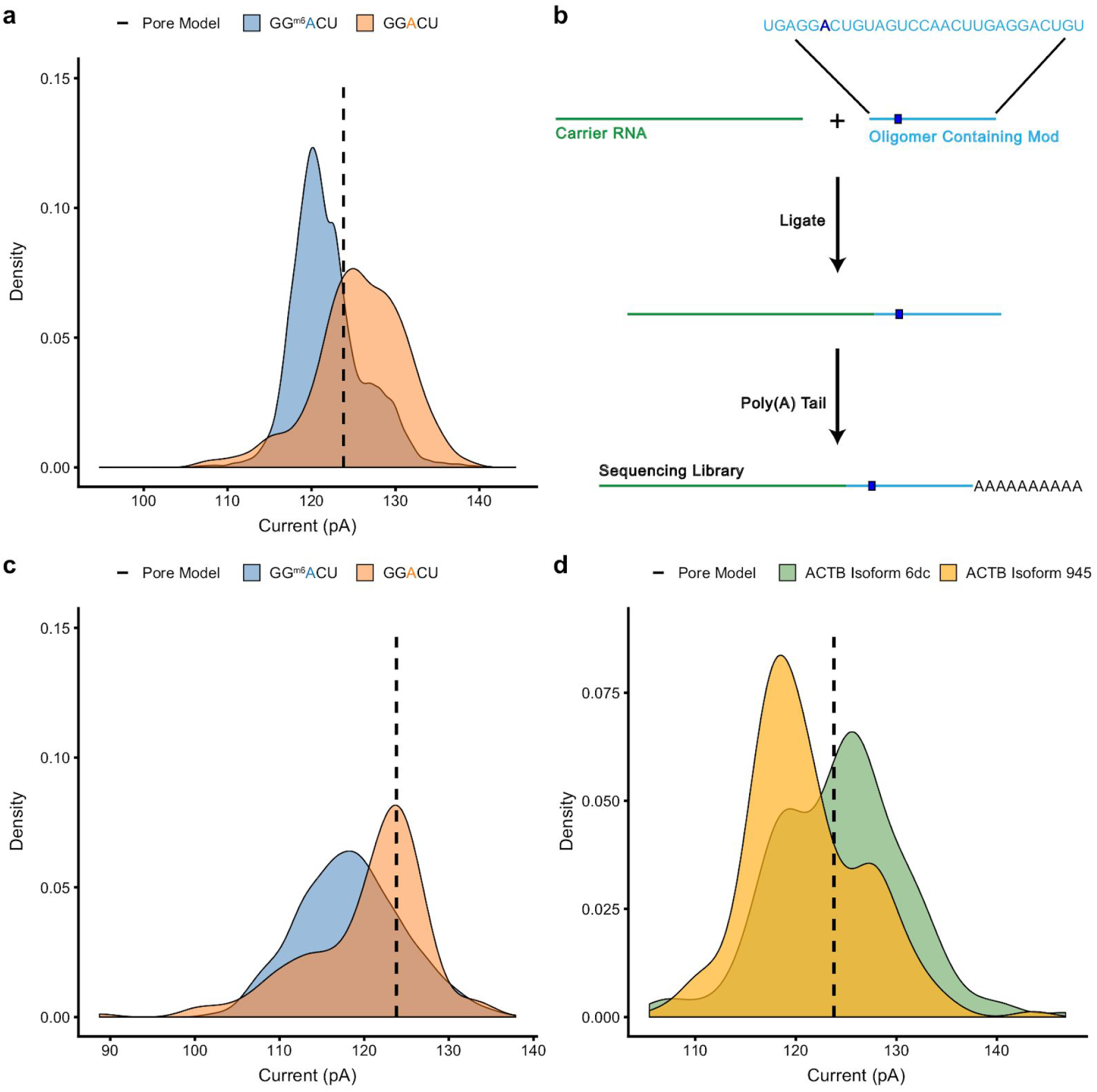
Nanopore detection of m6A base modifications. (a-c) The extent and directionality of current (pA) shift at GGACU motifs compared between our native RNA sequencing dataset, an *in vitro* transcribed dataset, and synthetically generated oligos drawn from a known highly methylated position within *EEF2* and the same position within unmodified *in vitro* transcribed GM12878 RNA. (a) Comparing current signal from m6A-modified and unmodified GGACU motifs in the native RNA dataset and *in vitro* transcribed dataset (b) Schematic for the oligomer-ligation preparation. A synthetic RNA oligomer (Trilink Biotechnologies) is ligated to a carrier RNA, followed by *in vitro* polyadenylation in preparation for sequencing. (c) Comparing current signal from m6A-modified and unmodified GGACU motifs from molecules sequenced using the assay described in (b). (d) Current distributions for GGACU motifs within actin (ACTB) gene isoforms.

To cross-validate this result, we designed and synthesized a 29 base oligomer containing both m6A and an unmodified adenosine within the GGACU METTL3 motif (Figure 7b). This short oligomer was ligated to a carrier sequence (*in vitro* transcribed firefly luciferase (NEB HiScribe) using T4 RNA ligase 1. The resulting product was polyadenylated and sequenced on the MinION. Using nanopolish eventalign, we extracted the current levels for the k-mers which contained the modified base and the unmodified base, revealing a clear difference (Figure 7c) consistent with the *EEF2* result.

To determine if m6A modifications differed between isoforms of the same gene, we screened isoform Set B for ionic current changes at the GGACU motif. We found 57 genes (158 isoforms) where the average current levels at GGACU differed more than 3 pA between gene isoforms with >100X positional coverage. An example is illustrated for *ACTB* (Figure 7d, **Supplementary Figure 3**). Interestingly, for *ACTB* both isoforms appeared to be modified at their respective GGACU motifs, but the ratio of modified to unmodified sequences differed.

Another post-transcriptional modification, A-to-I RNA editing^49^, commonly occurs in introns, UTRs, and Alu elements. It plays an important role in splicing and regulating innate immunity^50,51^ and is associated with epilepsy, autism, neurodegenerative diseases, and cancer^52^. NGS detects A-to-I editing as a nucleotide variant in cDNA sequences (A-to-G).

We found three lines of evidence for A-to-I edits in the *AHR* nanopore data (Figure 8). First, previous work on 16S rRNA documented the presence of systematic base miscalls in regions bearing modified RNA bases^19^, and indeed there are systematic base miscalls at putative inosine bearing positions in the *AHR* data (Figure 8a). Second, putative inosines were detected as a base change (A-to-G) in nanopore cDNA sequence data relative to the reference (single inosine for the CUACU kmer, and multiple inosines for the AAAAA kmer, Figure 8a). Third, we expected that ionic current distributions for these inosine containing kmers would be different from A containing kmers. Using nanopolish eventalign, we extracted the ionic current distributions for these putative inosine containing kmers and compared them to their canonical RNA counterparts. The ionic current distribution for the putative single inosine kmer (CUICU) showed substantial overlap with the canonical kmer CUACU (Figure 8b). The ionic current distribution for AAAAA kmer was more complex with at least 5 local maxima, likely reflecting multiple inosines (Figure 8c). The complexity of changes in ionic current distributions appeared to be consistent with the number of putative inosine containing positions.

**Figure 8.**
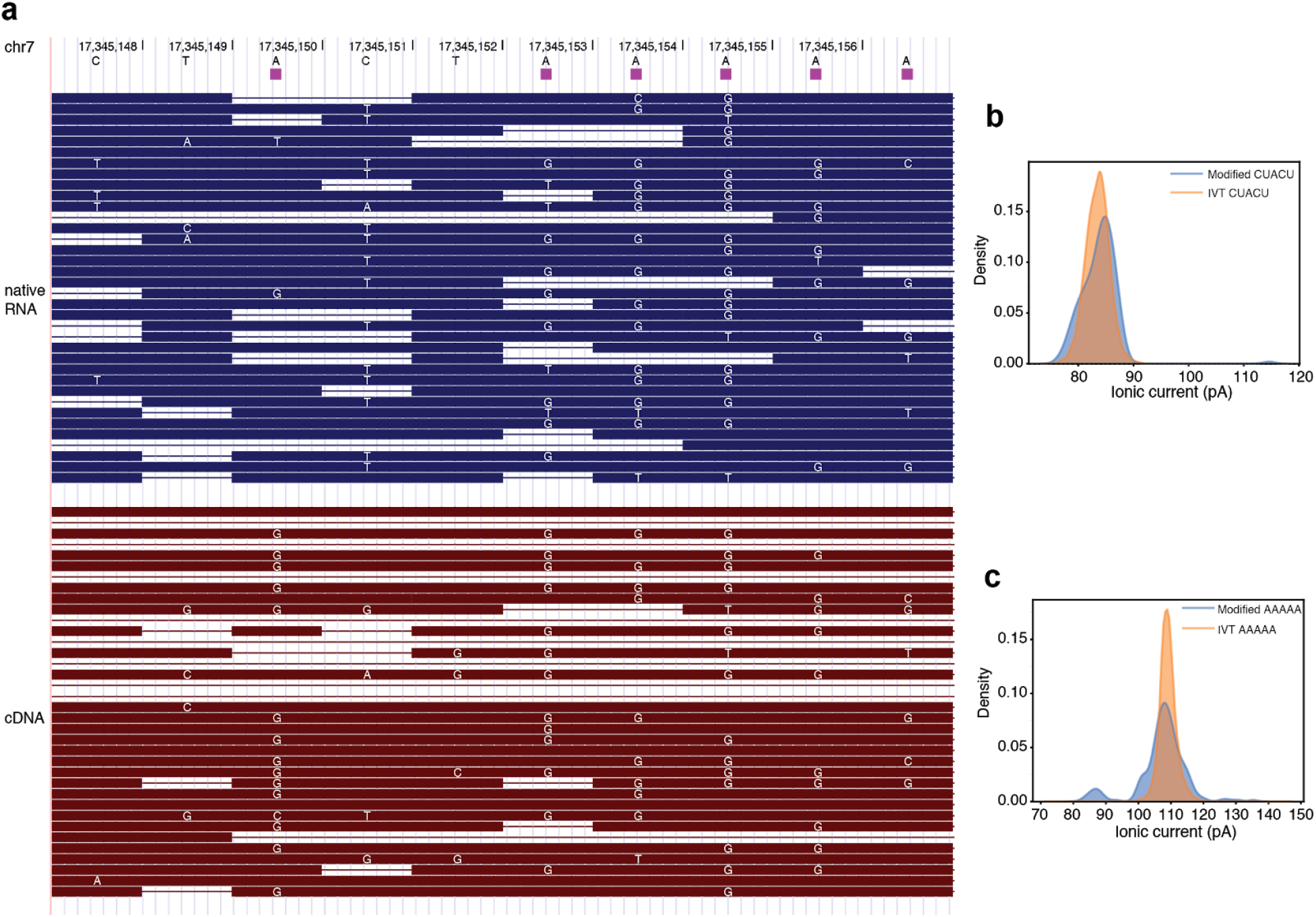
Nanopore detection of inosine base modifications. (a) Genome browser shot showing putative inosine positions as characterized by RADAR^78^ (magenta squares between putative A-to-I sites in the reference sequence and the alignment panel), and read alignments for nanopore native RNA (blue) and cDNA data (brown). Of note are base miscalls in native RNA data at or near A-to-I sites denoted by RADAR, and G calls at the corresponding positions in cDNA data. (b) Ionic current distributions for CUACU kmer from native RNA (blue) vs. IVT data (orange). (c) Ionic current distributions for AAAAA kmer from native RNA (blue) vs. IVT data (orange).

## DISCUSSION

Nanopore RNA sequencing has two important features: 1) The sequence composition of each strand is read as it existed in the cell. This permits direct detection of post-transcriptional modifications including base alterations and polyadenylation; and 2) reads can be continuous over many thousands of nucleotides. Although each of these features is useful in itself, the combination is unique and likely to provide new insights into RNA biology. The two principal drawbacks of the present ONT nanopore RNA sequencing platform are relatively higher error rates, and inability to read approximately 10-15 nucleotides at the 5′ end of each strand.

We were concerned that short fragment reads in our data might be caused by RNA degradation on the nanopore flow cells during typical 48-hour sequencing runs. To address this, we quantified the fraction of mitochondrial-encoded MT-CO1 transcripts that were full length as a function of experiment run time. We found minimal (~5%) reduction in the full length fraction over 48 hours. Preliminary analysis indicated that read truncations were more often caused by electronic signal noise associated with enzyme motor stalls during strand translocation, or by stray current spikes of unknown origin. Nonetheless, a substantial fraction of fragmented nanopore RNA reads appear to be biological in origin.

When combined with more accurate short Illumina reads, long nanopore reads allowed for end-to-end documentation of RNA transcripts bearing numerous splice junctions, which would not be possible using either platform alone. Using our most stringent filters, 65.3% of detected isoforms (over 30,000 total) from the nanopore GM12878 transcriptome were not annotated in GENCODE v24. Of the 65.3% novel isoforms, 25.5% (8,514) have retained introns, 25.2% (8,407) contain new combinations of annotated splice junctions, 9.2% (3,079) contain completely novel exons, and 0.7% (229) are at putative novel gene loci (**Supplementary Figure 4**). Other long-read transcriptome sequencing studies have observed similar numbers of novel isoforms (e.g., 35.6% and 49%)^53,54^. While many of these novel isoforms are low abundance and their protein coding potential unknown, transcript variation is still important to catalogue as subtle splicing changes can impact function^55,56^. It is also noteworthy that the number of detected isoforms did not saturate using the nanopore poly(A) RNA dataset (Figure 4b) indicating that greater sequence depth will be necessary to give a comprehensive picture of the GM12878 poly(A) transcriptome.

A variety of techniques have been used to examine allele-specific expression (ASE)^57-63^. This has proven to be clinically useful because ASE is frequently increased in human cancers as a result of copy number variations^60^. However, the ability to identify instances of allele-specific expression (ASE) is limited by the requirement of a heterozygous variant within the sequencing read. Further, the results of ASE analysis can be difficult to interpret, as it remains unclear when ASE represents a pathology versus normal biology. Biased expression from one allele can occur stochastically as part of normal physiology, and can change at different times in the cell cycle^64^. A targeted study of transformed cells from 69 individuals found expression to be allele-biased at 88% percent of the genes evaluated^65^, and suggested that underlying cis-regulatory transcriptional activation mechanisms enhanced this phenomena.

Our nanopore approach for ASE discovery has the advantage of long reads, but the disadvantage of high base call errors. We attempted to mitigate the effects of nanopore sequencing errors by imposing stringent thresholds for assigning reads to allele and a lower FDR during ASE analysis. The number of genes that we reported as demonstrating ASE (492) is likely an underestimation. Error correction to assign the read where parental allele is ambiguous as well as deeper coverage of low abundance reads would likely reveal further subtleties of ASE^65^ and increase the number of genes where ASE is significant. Of the significant ASE genes in our dataset, the X chromosome provides a good gene set to evaluate allele-specific expression because of its known single-allele inactivation for dosage compensation in females. We report near exclusive use of the maternal X-chromosome, with the only paternal reads originating from the Xist locus, consistent with previous findings^57^.

Polyadenylation of RNA 3′ ends regulates RNA stability and translation efficiency by modulating RNA-protein binding and RNA structure^66^. However, transcriptome-wide poly(A) analysis has been difficult because standard Illumina-based measurements are impacted by basecalling and dephasing errors^67^. Recently implemented modifications of the Illumina strategy address these limitations^67,68,69^; but do not resolve more complex questions, e.g. relationships between splicing and poly(A) length. Nanopore poly(A) tail length estimation using nanopolish-polya offers several advantages: i) Each RNA strand is read directly so errors due to priming of internal poly(A) segments cannot occur; ii) the entire length of a transcript can be read; therefore, linkage between a particular isoform, its modifications, and poly(A) tail length can be documented directly; and iii) no additional preparative steps are required because the ionic current segment associated with the 3′ poly(A) tail is part of the overall nanopore sequencing signal. However, currently native RNA sequencing of poly(A) tails will miss a portion of mRNA species. For example, mRNAs with short (<10) poly(A) tails or with uridylation stretches would not be efficiently hybridized by the splinted poly(T) sequencing adapter. Alternative library preparations will need to be employed to resolve this issue.

Our preliminary studies revealed differences in poly(A) length distribution between different genes, and between different isoforms of the same gene. We note in particular a difference in poly(A) tail length between spliced transcripts and unspliced transcripts bearing introns. This suggests that time course experiments investigating RNA processing and decay kinetics^70^ could be possible with this technology. This might be especially useful because deadenylation is a core part of the RNA degradation pathway^71^.

We have demonstrated detection of N6-methyladenosine and inosine modifications in human poly(A) RNA. This validates prior work^18,19^ which showed modification-dependent ionic current shifts associated with m6A (*S. cerevisiae*)^18,19^ and m7G (*E. coli*)^18,19^. Differences in m6A modification level proved to be discernible at the isoform level for human ACTB mRNA (Figure 7d), thus documenting splicing variation and modification changes simultaneously.

Although other methods exist for high throughput analysis of RNA modifications^72^, they often require enrichment which limits quantification, and they are usually short-read based. The latter precludes analysis of long-distance interactions between modifications, and between modifications and other RNA features such as splicing and poly(A) tail length. The capacity to detect these long-range interactions is likely to be important given recent work suggesting links between RNA modifications, splicing regulation, and RNA transport and lifetime^73,74^. We argue that nanopore native RNA sequencing could deliver this long-range information for entire transcriptomes. However, this will require algorithms trained on large, rigorously cross-validated datasets as has been accomplished for cytosine modifications in genomic DNA^75,76^.

## CONCLUSIONS

Oxford Nanopore devices sequence long native RNA strands directly. In this study, we showed that these long reads improved human poly(A) RNA isoform characterization, including allele specificity. Because native RNA strands were read directly, m6A and inosine nucleotide modifications could be detected absent intermediate preparative steps. We introduced a new tool (nanopolish-polya) that estimates 3′ poly(A) tails on individual RNA strands based on nanopore ionic current signals. Applied to the GM12878 transcriptome, it revealed differences in RNA poly(A) tail lengths between genes, and between isoforms of the same gene.

## ONLINE METHODS

### GM12878 cell tissue culture

GM12878 cells (passage 4) were received from the Coriell Institute and cultured in RPMI media (Invitrogen cat# 21870076) supplemented with 15% non heat-inactivated FBS (Lifetech cat# 12483020) and 2mM L-Glutamax (Lifetech cat# 35050061). Cells were grown to a density of 1 × 10^6^ / ml before subsequent dilution of ⅓ every ~3 days, and expanded to 9 x T75 flasks (45 ml of media in each). Cells were centrifuged for 10 min at 100 x g (4°C), washed in 1/10th volume of PBS (pH 7.4) and combined for homogeneity. The cells were then evenly split between 8 × 15ml tubes and pelleted at 100g for 10 mins at 4°C. The cell pellets were then snap frozen in liquid Nitrogen and immediately stored at -80°C before shipping on dry ice. Two tubes of 5 × 10^7^ frozen GM12878 cell pellets from passage 10 from a single passage, cultured at UBC, were distributed and used at UBC, OICR, JHU and UCSC. Two tubes of cells from passage 11 was distributed to UoN from UBC, and an independently cultured passage of GM12878 was used at UoB. (University of British Columbia (UBC), University of Birmingham (UoB), Ontario Institute of Cancer Research (OICR), Johns Hopkins University (JHU), University of Nottingham (UoN), and University of California Santa Cruz (UCSC))

### Total RNA Isolation

The following protocol was used by each of the six institutions. Four ml of TRI-Reagent (Invitrogen AM9738) was added to a frozen pellet of 5 × 10^7^ GM12878 cells and vortexed immediately. This sample was incubated at room temperature for 5 minutes. Four hundred μl BCP (1-Bromo-3-chloro-propane) or 200 μl CHCl3 (Chloroform) was added per ml of sample, vortexed, incubated at room temperature for 5 minutes, vortexed again, and centrifuged for 10 minutes at 12,000g (4°C). The aqueous phase was pooled in a LoBind Eppendorf tube and combined with an equal volume of isopropanol. The tube was mixed, incubated at room temperature for 15 minutes, and centrifuged for 15 minutes at 12,000g (4°C). The supernatant was removed, the RNA pellet washed with 750 μl 80% ethanol and centrifuged for 5 minutes at 12,000g (4°C). The supernatant was removed. The pellet was air-dried for 10 minutes, resuspended in Nuclease free water (100 μl final volume), quantified, and either stored at -80°C or processed further by poly-A purification.

### Poly(A) RNA isolation

One hundred μg aliquots of total RNA were diluted in 100 μl of nuclease free water and poly-A selected using NEXTflex Poly(A) Beads (BIOO Scientific Cat#NOVA-512980). Resulting poly-A RNA was eluted in Nuclease free water and stored at -80°C.

### MinION native RNA sequencing of GM12878 poly-A RNA

Biological poly-A RNA (500-775 ng) and a synthetic control (Lexogen SIRV Set 3, 5 ng) were prepared for nanopore direct RNA sequencing generally following the ONT SQK-RNA001 kit protocol, including the optional reverse transcription step recommended by ONT. One difference from the standard ONT protocol was use of Superscript IV (Thermo Fisher) for reverse transcription.

RNA sequencing on the MinION and GridION platforms was performed using ONT R9.4 flow cells and the standard MinKNOW protocol script

(NC_48Hr_sequencing_FLO-MIN106_SQK-RNA001)

recommended by ONT, with one exception. We restarted the sequencing runs at several time points to improve active pore counts and throughput during the first 24hrs.

### cDNA synthesis

First strand cDNA synthesis was performed using Superscript IV (Thermo Fisher) and 100 ng of poly-A purified RNA combined with 0.5 ng of the SIRV set 3 control. Reverse transcription and strand-switching primers were provided by ONT in the SQK-PCS108 kit. After reverse transcription, PCR was performed using LongAmp Taq Master Mix (NEB) under the following conditions: 95°C for 30 seconds, 11-15 cycles (95°C for 15 seconds, 62°C for 15 seconds, 65°C for 15 minutes), 65°C for 15 minutes, hold at 4°C. The 15 cycle PCR was performed when using the SQK-PCS108 kit and 11 cycle PCR was performed when using the SQK-LSK308 kit. PCR products were purified using 0.8X AMPure XP beads.

### MinION sequencing of GM12878 cDNA

cDNA sequencing libraries were prepared using 1 μg of cDNA following the standard ONT protocol for SQK-PCS108 (1D sequencing) or SQK-LSK308 (1D^2 sequencing) with one exception. That is, we used 0.8X AMPure XP beads for cleanup. We used standard ONT MinKNOW scripts for MinION sequencing with one exception. That is, we restarted the sequencing runs at several time points to improve active pore counts and throughput during the first 24 hours.

### Acquiring continuous data for nanopore sequencing runs

For a subset of runs “bulk files” containing continuous raw current traces and read decisions made by MinKNOW were recorded for more detailed analysis. This can be enabled in MinKNOW by looking at “Additional options” under “Output” when configuring a run to start in MinKNOW. Options were set to capture Raw signal data and the Read Table. Events were not captured to reduce file size^79^.

### Length analysis of mitochondrial protein-coding transcripts

In this analysis, we limited the test population for each gene to reads that aligned to a 50 nt sequence at the 3′ prime end of its ORF, except for MT-ND5 where alignment was to a 50 nt sequence at the end of its 568 nt 3′ UTR. Full length was defined as extending to at least within 25 nt of the genes expected 5′ terminus. This limit was chosen because the processive enzyme that regulates RNA translocation is distal from the CsgG nanopore limiting aperture and necessarily falls off before the 5′ end is read. The sharpest coverage drop-off is typically at 10 nt from the 5′ transcript end; we chose the 25 nt limit to ensure that all likely full length reads were captured in the count.

### *In vitro* transcription

First strand cDNA synthesis was performed using Superscript IV (Thermo Fisher) and 100 ng of poly-A purified RNA combined with 0.5 ng of the SIRV set 3 control. Reverse transcription and strand-switching primers were provided by ONT in the SQK-PCS108 kit. After reverse transcription, PCR was performed using LongAmp Taq Master Mix (NEB) under the following conditions: 95°C for 30 seconds, 11-15 cycles (95°C for 15 seconds, 62°C for 15 seconds, 65°C for 15 minutes), 65°C for 15 minutes, hold at 4°C. An 11 cycle PCR was performed using recommendations from the SQK-LSK308 kit and a modified version of the primer that included a the priming site for T7 RNA polymerase as recommended by NEB at

(https://www.neb.com/products/e2040-hiscribe-t7-high-yield-rna-synthesis-kit#Product%20Information).

PCR products were purified using 0.8X AMPure XP beads. Next, in vitro transcription was performed using the NEB HiScribe T7 High Yield RNA Synthesis Kit. The IVT product was also further poly(A) tailed using the same kit. The resulting IVT RNA was purified using LiCl precipitation and then adapted for RNA sequencing on the MinION.

#### Analysis pipelines

##### Basecalling, alignments, and percent identity calculations

We used the ONT Albacore workflow (version 2.1.0) for basecalling direct RNA and cDNA data. A tunable filter in Albacore then classifies individual strand reads as pass or fail based-on their sequence quality. We used minimap2^80^ (recommended parameters) to align the nanopore RNA and cDNA reads to the GRCh38 human genome reference^81^ and to the GENcode transcriptome reference^82^. We used marginStats^83^ to calculate alignment identities and errors for pass RNA strand reads and pass 1D cDNA strand reads. Read identities were also calculated for a number of the plots generated using the nanopore scripts provided by Aaron Quinlan^77^. Substitutions were calculated using custom scripts available within marginAlign^26^.

### Kmer analysis

We used the high confidence isoforms yielded by FLAIR as reference sequences. For kmer analysis, we aligned the native RNA and cDNA reads to these isoforms and created a map of read to reference sequences. We calculated expected kmer counts from the set of reference sequences and observed kmer counts from the set of read sequences. For plotting purposes, we normalized the read and reference counts to coverage per megabase. The scripts are available within marginAlign^26^.

### Isoform detection and characterization

To define isoforms from the sets of native RNA and cDNA reads, we used FLAIR v1.1, a version of FLAIR^25^ with additional considerations for native RNA nanopore data. From our analysis, we first removed reads generated from the lowest quality flowcell run as many of those reads were considered to potentially be truncated. Next, we aligned pass reads to the genome with minimap2, retaining only primary alignments. Using FLAIR-correct, we corrected the genome-aligned reads providing splice site evidence from Gencode v24 annotations and Illumina short-read sequencing of GM12878. This step also removes reads containing splice junctions not present in the annotation or short-read data. We then filtered for reads with TSSs falling within promoter regions derived from an HMM based on ENCODE ChIP-Seq data of nine factors ^34,35^ . The corrected and filtered reads were then given to FLAIR-collapse to produce a nanopore-specific reference containing high-confidence isoforms with at least 5 supporting reads (set A). In short, FLAIR-collapse first generates a first-pass isoform set based on reads grouped by the splice junctions they contain. The first-pass set contains isoforms matching annotations as well as isoforms specifically expressed in native RNA reads, capturing nanopore-exclusive isoforms. The final isoform set is created by taking all pass reads, including those that were filtered out previously, and realigning to the first-pass isoform set. Additionally, to produce a more stringent set of isoforms (set B), any isoforms that were a subset of a longer isoform were filtered out.

Productivity was assessed according to an NMD rule where if a premature termination codon is located 55 nt or more upstream of the last exon-exon junction the transcript is considered unproductive^84^. Genes that did not contain an annotated start codon were considered noncoding genes. To account for differences in expression between coding and noncoding genes, the number of isoforms per gene was normalized by dividing by the (number of reads aligned to the gene/100).

Novel isoforms were defined as those with a unique splice junction chain not found in annotations. Isoforms were considered intron-retaining if they contained an exon which spanned another isoform’s splice junction completely. Novel isoforms with novel exons were defined as isoforms with at least one exon that does not overlap any existing annotated exons. Isoforms at novel loci were defined as isoforms that only contain novel exons.

### Defining promoter regions in GM12878 for isoform filtering

Promoter chromatin states for GM12878 were downloaded from the UCSC Genome Browser in BED format from the hg18 genome reference. Chromatin states were derived from an HMM based on ENCODE ChIP-Seq data of nine factors^34,35^. The liftover tool^85^ was used to convert hg18 coordinates to hg38. The active, weak, and poised promoter states were used.

### Haplotype Assignment and Allele-Specific Analysis

We obtained genotype information for GM12878 from existing phased illumina platinum genome data generated by deep sequencing of the cell donors’ familial trio^86^. The bcftools package was used to filter for only variants that are heterozygous in GM12878. Starting with aligned reads, we used the extractHAIRS utility of the haplotype-sensitive assembler HapCUT2^38^ to identify reads with allele-informative variants. For stringent analysis, we required a read to contain at least two variants, and required that greater than 75% of identified variants agreed on the parental allele of origin -- this stringent threshold was selected to reduce the chances of a false positive from nanopore sequencing errors. Through this approach each read was annotated as maternal, paternal or unassigned. We used the output of the FLAIR pipeline for annotating the gene for each of the reads. To identify genes which demonstrated a very strong bias for a single allele, we performed a binomial test of all reads assigned to parental allele, with a FDR of 0.01. We also visually inspected numerous genes displaying genes demonstrating allele-specificity using IGV, to increase our confidence in proper mapping of the reads and evaluate the presence of variants.

We further integrated this haplotype-specific analysis with our isoform pipeline to explore for the presence of allele-specific isoforms. If greater than 80% of reads for a specific isoform originated from a single parental allele, the isoform was assigned as allele specific. We then filtered for any genes which contained both maternal and paternal allele-specific isoforms, and visually inspected these isoforms using IGV to compare location of variants and splicing events.

### Poly(A) tail length analysis

See **Supplemental Note** describing the operation of nanopolish-polya for estimation of polyadenylated tail lengths from native RNA sequencing. We applied this tool to our dataset, focussing on primary aligned reads. We then correlated the called poly(A) length per read with which FLAIR detected isoform it aligned to. First, the distribution of reads which aligned to nuclear transcripts was compared to reads aligned to mitochondrial transcripts. We then categorized genes according to the number of reads associated with them ([5-1,000], [1,000-20,000], and [20,000-] reads). For each category, genes were ranked according to their mean poly(A) tail length. We examined the two longest, two shortest and middle length of this list. Genes with a complex distribution were further investigated further at the isoform level, examining only the isoforms which had at least 20 reads supporting them. Finally, isoforms were annotated as spliced, or intron retained from the FLAIR labels, and the distribution of average poly(A) for transcripts belonging to these two categories plotted.

### Modification detection and analysis

In order to detect putatively modified sites within our native RNA dataset, we first determined transcripts likely to be methylated and looked for regions of difference relative to the expected model distribution and in-vitro transcribed (IVT) data. We focused our initial efforts on the m6A modification, which is written by the METTL3-METTL14-WTAP complex within the consensus motif DRACH. By intersecting native RNA datasets with transcripts enriched in previous m6A immunoprecipitation pulldowns of human cell lines ^47,48^, we identified several candidate genes to examine. Alignments of reads pertaining to candidate genes in both native and IVT RNA datasets were fed into the eventalign module of nanopolish ^87^, which allows for the alignment of electrical events to reference k-mers, with the output of this module being the event level mean (current, pA) and standard deviation by kmer of the input reads ^23^. This workflow then allows us to compare, for example, the current at GGACU within the 3′ UTR of the eEF2 gene in native RNA, to the same k-mer at the same position for in-vitro transcribed RNA. The expectation is that a modification within the kmer of interest in native RNA would shift the current measured at that position, differentiating it from that of the IVT alignments.

The extent and directionality of current shift observed by m6A modification within the GGACU motif was orthogonally investigated using an in-vitro oligomer ligation assay, preparation methods described above, and current within the modified and unmodified GGACU motifs within the synthetic oligomer were compared using eventalign.

For detecting A-to-I editing in the data, we identified a candidate gene that had previously been shown to be inosine rich and then analyzed the alignments for native RNA and cDNA data to the human genome. We focused our efforts at the human aryl hydrocarbon receptor (AHR) gene. We analyzed the alignments in the 3′-UTR region initially using the genome browser and then nanopolish eventalign. Using the genome browser, we searched for base variant calls in cDNA data and systematic base miscalls in native RNA data. We then used nanopolish eventalign to yield ionic current distributions for inosine-containing kmers and compared them with distributions for their respective canonical kmers arising from whole GM12878 poly-A RNA IVT nanopore data. We used IVT data from chromosome 7 for the comparison.

## ACKNOWLEDGEMENTS

The authors are grateful for support from the following individuals. Libby Snell, Botond Sipos and Dan Turner (ONT) provided materials and advice relevant to the 3′ poly(A) standards used to test nanopolish-polya. Daniel Garalde (ONT) provided early advice on use of the MinION for RNA sequencing. Nicholas Conrad gave insight into the correlation of intron retention and poly(A) tail length. Mark Diekhans reviewed the isoform analysis. The authors thank Andrew Beggs, Louise Tee and Tom Nieto (University of Birmingham, UK) for providing cell cultures used in the Birmingham sequencing runs. The project was supported by the following grants: NIH HG010053 (AB, BP, & MA), NIH 5T32HG008345 (AT), NIH HG009190 (WT, JTS), NIH U54HG007990 (BP), U01 HL137183-02 (BP), Oxford Nanopore Research Grant SC20130149 (MA), National Institutes of Health Research Surgical Reconstruction and Microbiology Research Centre (JQ), Medical Research Council CLIMB Fellowship (NL), Wellcome Trust 204843/Z/16/Z (ML), BBSRC BB/N017099/1 and BB/M020061/1 (ML), the Canada Research Chair in Biotechnology and Genomics-Neurobiology (TPS), the Canadian Institutes of Health Research (#10677; TPS), the Canadian Epigenetics, Environment and Health Research Consortium (TPS), the Koerner Foundation (TPS), the Ontario Institute for Cancer Research through funds provided by the Government of Ontario (JTS).

